# Toward Minimally Invasive Therapeutic Ultrasound: Ultrasound-guided Ablation in Neuro-oncology

**DOI:** 10.1101/2020.04.25.061788

**Authors:** Micah Belzberg, Smruti Mahapatra, Francisco Chavez, Kyle Morrison, Kah Timothy Xiong, Nao J. Gamo, Stephen Restaino, Rajiv Iyer, Mari Groves, Nitish Thakor, Nicholas Theodore, Mark G. Luciano, Henry Brem, Alan R. Cohen, Amir Manbachi

**Author notes:** **Corresponding Author’s Address:** Amir Manbachi, Ph.D., Departments of Neurosurgery and Biomedical Engineering, Johns Hopkins University, 600 N. Wolfe St., Meyer 8-181C, Baltimore, MD 21287, USA. **Funding:** This study was funded by Maryland Technology Development Corporation’s Maryland Innovation Initiative, Coulter Foundation, National Science Foundation’s I-Corps program and Johns Hopkins University, Whiting School of Engineering’s Cohen Translational Funding opportunities. **Financial Disclosures:** Kyle Morrison, Francisco Chavez, and Kah Timothy Xiong are employees of Sonic Concepts, Inc. Dr. Brem is a consultant serving as Medical Advisory Board Chairman for InSightec. He is also on the board of directors for Galen Robotics. Dr. Nao J Gamo is the founder and CEO of Neurosonics Medical, Inc. Dr. Stephen Restaino is the Director of Engineering at Maryland Development Center, a startup studio supporting local medical device innovations.

## Abstract

**Introduction:** To improve patient outcomes (eg, reducing blood loss and infection), practitioners have gravitated toward noninvasive and minimally invasive surgeries (MIS), which demand specialized toolkits. Focused ultrasound, for example, facilitates thermal ablation from a distance, thereby reducing injury to surrounding tissue. Focused ultrasound can often be performed noninvasively; however, it is more difficult to carry out in neuro-oncological tumors, as ultrasound is dramatically attenuated while propagating through the skull. This shortcoming has prompted exploration of MIS options for intracranial placement of focused ultrasound probes, such as within the BrainPath^™^ (NICO Corporation, Indianapolis, IN). Herein, we present the design, development, and *in vitro* testing of an image-guided, focused ultrasound prototype designed for use in MIS procedures. This probe can ablate neuro-oncological lesions despite its small size.

**Materials & Methods:** Preliminary prototypes were iteratively designed, built, and tested. The final prototype consisted of three 8-mm-diameter therapeutic elements guided by an imaging probe. Probe functionality was validated on a series of tissue-mimicking phantoms.

**Results:** Lesions were created in tissue-mimicking phantoms with average dimensions of 2.5×1.2×6.5mm and 3.4×3.25×9.36mm after 10- and 30-second sonification, respectively. 30s sonification with 118W power at 50% duty cycle generated a peak temperature of 68°C. Each ablation was visualized in real time by the built-in imaging probe.

**Conclusion:** We developed and validated an ultrasound-guided focused ultrasound probe for use in MIS procedures. The dimensional constraints of the prototype were designed to reflect those of BrainPath trocars, which are MIS tools used to create atraumatic access to deep-seated brain pathologies.

**HIGHLIGHTS:**

- An ultrasound-guided, focused ultrasound prototype was developed and validated
- The therapeutic transducer (1.5MHz) consisted of three 8-mm circular elements
- Elements were placed on a 9×32mm curved rectangular aperture: 45mm radius curvature
- Functionality was examined on tissue-mimicking phantoms
- 2.5×1.2×6.5mm and 3.4×3.25×9.36mm lesions were seen for 10 and 30s sonification

## 1. INTRODUCTION

Focused ultrasound is an appealing tool for use in both noninvasive and minimally invasive surgical (MIS) procedures, as it can ablate pathologic tissue from a distance.^1–3^ This is particularly helpful in patients diagnosed with inoperable tumors.^4,5^ In neurosurgery, noninvasive (transcranial) focused ultrasound helmet systems have been successfully used to treat movement disorders, including Parkinson disease and essential tremor.^6^ However, their use for ablation of neuro-oncological tumors faces certain drawbacks;^3^ namely, significant attenuation of the ultrasonic wave during propagation in the skull, which requires high ultrasonic powers directed at the patient’s head.^7–9^ In response to these drawbacks, MIS focused ultrasound approaches have been contemplated, as they may require only miniature probes to ablate intracranial lesions from within the cranium.^10,11^

One such example is a minimally invasive, focused ultrasound probe capable of being placed within the BrainPath™ (NICO Corporation, Indianapolis, IN), a device that allows atraumatic access to brain oncology (eg, deep-seated tumors) in MIS settings (**Fig. 1**). The development of products like BrainPath™ represents a growing trend toward MIS options in healthcare, where surgical interventions are increasingly being performed with minimally invasive approaches or keyhole procedures to enhance patient outcomes and reduce blood loss or infection.^12^ Of course, this trend requires development of novel, precise, miniature surgical instruments. Although larger transducer surface areas allow for better energy deposition, MIS encourages increasingly smaller probes to reduce incision sizes and dissection requirements—and this tradeoff demands better understanding of acoustic designs to balance both criteria.^13,14,15^ In this study, we investigated preliminary focused ultrasound configurations for use in MIS settings, particularly for use in ablation of neuro-oncological lesions. Here, we report the development and *in vitro* testing of an initial series of probe designs small enough for use in MIS, yet with a large enough transducer surface area to generate the focal point ablations.

**Fig. 1.**
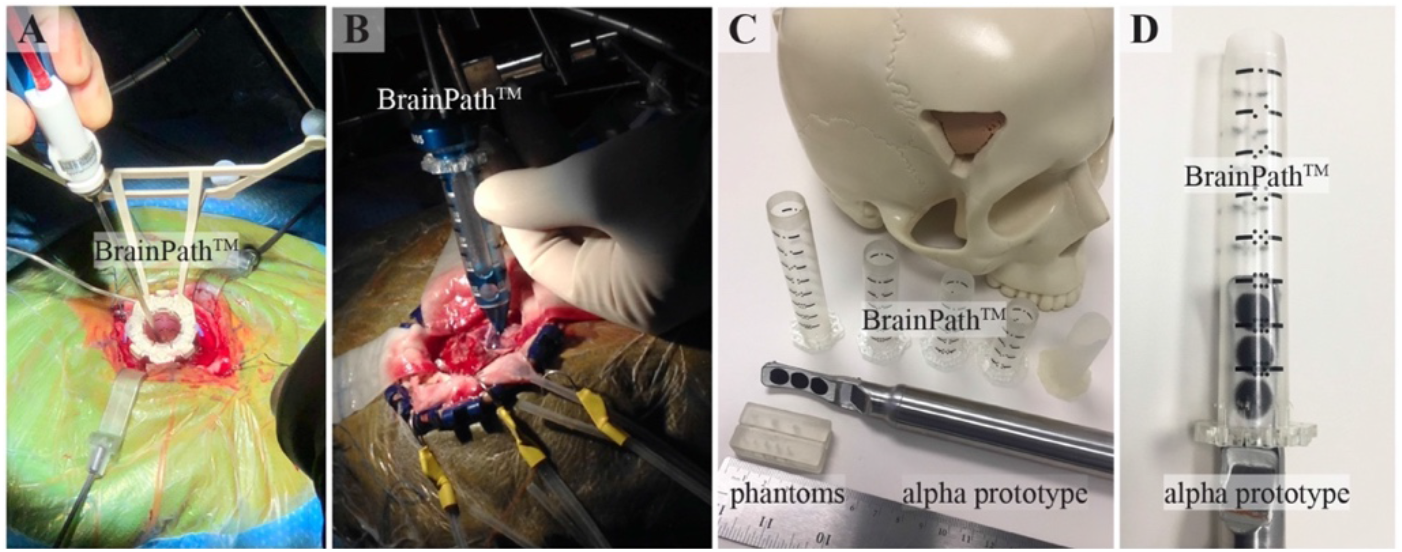
The BrainPath^™^ is a minimally invasive neurosurgical toolkit that can accommodate neuroendoscopic devices. (A&B) The BrainPath^™^ trocar is inserted into brain tissue to create atraumatic surgical access to deep-seated lesions. (C&D) The ultrasound-guided focused ultrasound alpha prototype probe described in this study is shown in relation to the BrainPath^™^ dimensions. *A and B contain original images presented with permission from NICO Corporation*.

## 2. MATERIALS AND METHODS

### Therapeutic Elements

Based on the dimensions of BrainPath™ and the guidelines for MIS and neuroendoscopy techniques (ie, burr holes measuring 18mm or smaller), the focused ultrasound transducer was designed to fit within a rectangular aperture of 9×32mm (**Figs. 1-2**).^16–18^ In order to accommodate the anatomy of a typical adult brain, where a tumor may be 4 to 5cm away from the probe placement within the BrainPath™, a transducer with a 45-mm radius of curvature (RoC) was designed to target a natural focal point lesion 4 to 5cm away. Building upon our prior simulation studies that investigated the effects of variations in transducer frequency, RoC, and power on the thermal dose and energy deposition in tissue, we developed and manufactured 2 alpha prototype transducer designs with center frequency of 1.5MHz (full width half maximum bandwidth: 1.20-1.80MHz) for *in vitro* validation (**Fig. 2**).^19,20^ Design I was a 1-piece, 9×32mm, cylindrically curved rectangular aperture with 45mm RoC. Design II contained three 8-mm-diameter circular elements placed on the curved geometry described above. The therapeutic array had a width of 12mm casing, which widened to 15mm to accommodate a built-in imaging array, as described below.

**Fig. 2.**
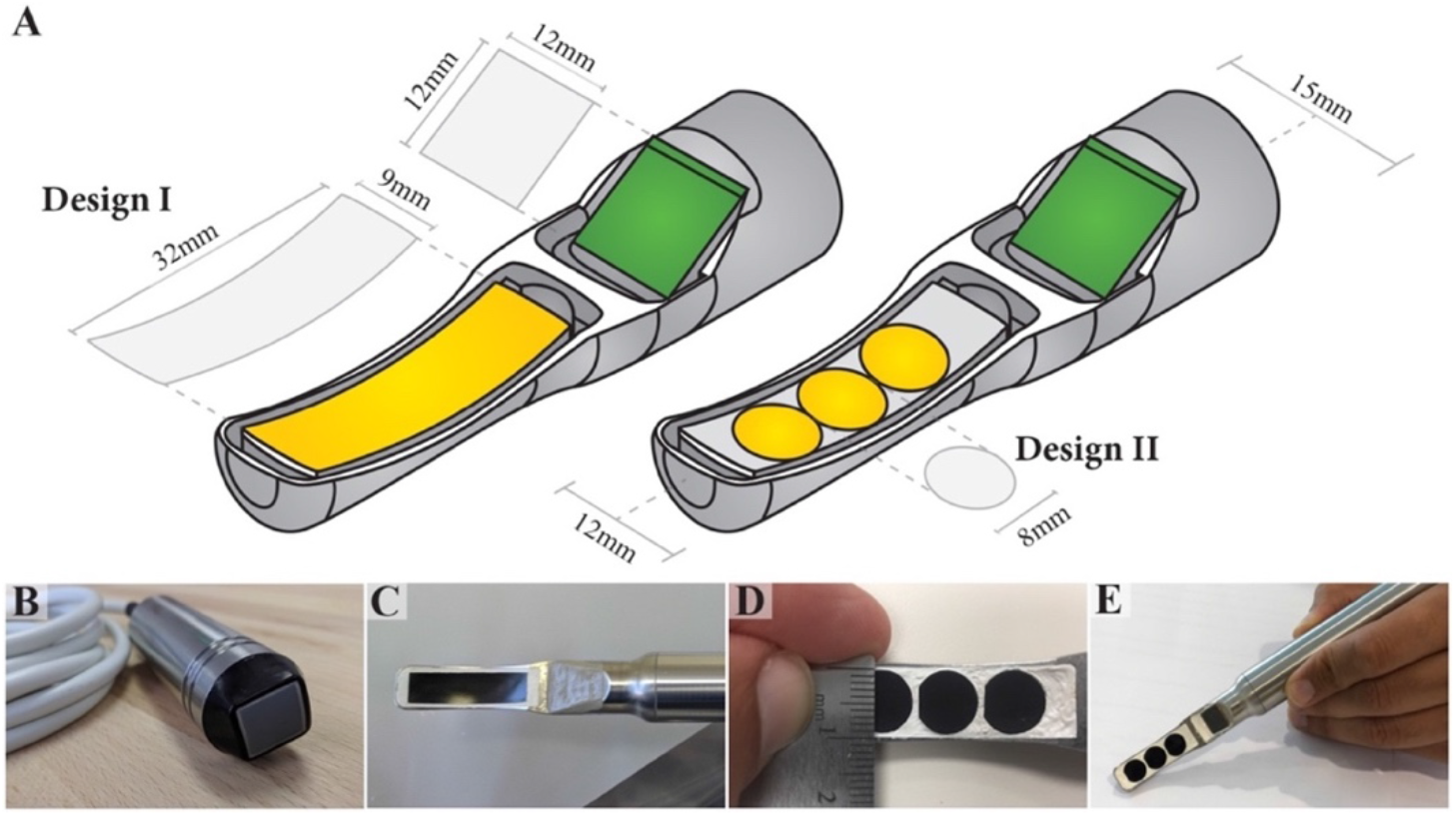
Probe designs and prototyping details. (A) Two late-stage designs of the custom probe containing a 12×12-mm imaging transducer adjacent to a therapeutic component. Design I is a 1-piece, 9×32mm, cylindrically curved rectangular aperture with 45mm RoC. Design II contains 3 circular therapeutic elements measuring 8mm in diameter, arrayed along a 9×32mm, cylindrically curved rectangular aperture with 45mm RoC. The therapeutic array fits within a casing 12mm in diameter, which widened to 15mm to accommodate the built-in imaging array. (B) A commercial imaging probe (Sonic Concepts IP-105, center frequency: 5MHz) which was integrated into our custom ultrasound-guided probe; (C) functional prototype of the custom probe (Design I) housed in a stainless steel casing; (D) 8-mm-diameter circular therapeutic elements (Design II); (E) complete prototype of Design II, containing both the imaging and therapeutic components.

### Imaging Elements

To reduce the need for intraoperative magnetic resonance imaging (MRI) guidance, a built-in ultrasound imaging probe was designed to demonstrate proof-of-concept for the ultrasound-guided focused ultrasound (USgFUS) approach studied here. An “off-the-shelf” IP-105 linear imaging probe from Sonic Concepts (Bothell, WA) was chosen. This 64-element, one-dimensional phased array (center frequency: 5.0MHz) was placed within the device housing and was tilted 30° to provide real-time visualization of the ablation lesion. Customized software was developed to drive the probe and to store, study, and modify signals. This software was developed on MATLAB (MathWorks Inc, Natick, MA) and installed on a Vantage 64 LE system (Verasonics Inc, Kirkland, WA).

### Experimental Assessment

*In vitro* testing of the preliminary prototype was performed by submerging the device in a 4-gallon test tank filled with tap water. Water was degassed to reduce the amount of dissolved gasses within the tank. Water temperature throughout experimental testing was 23°C, with less than 1°C variation. An acoustic absorber was placed inside the tank to reduce acoustic reflections from the tank edges. A calibrated oscilloscope as well as voltage and current probes were used to measure the voltage, current, and subsequently the net power of the device under test conditions. Using a three-dimensional (3D) printed holder created on a commercial 3D printer (Objet260, Connex3, Stratasys Ltd., Eden Prairie, MN), blocks of solid water phantoms were positioned at the focal point. Solid water is a tissue-mimicking phantom, proprietary to Sonic Concepts, with acoustic properties (ie, attenuation and propagation velocity) similar to those of water. The absorption coefficient at 1 MHz is ~0.01 dB/cm. Sound speed is 1,500 m/s. The phantoms were used to create predictable lesions and correlate results with calibrated hydrophone measurements. Lesion generation was observed both visually and via the built-in imaging probe driven by a Vantage 64 LE system (Verasonics, Inc., Kirkland, WA). The experimental design is shown in **Fig. 3.**

**Fig. 3.**
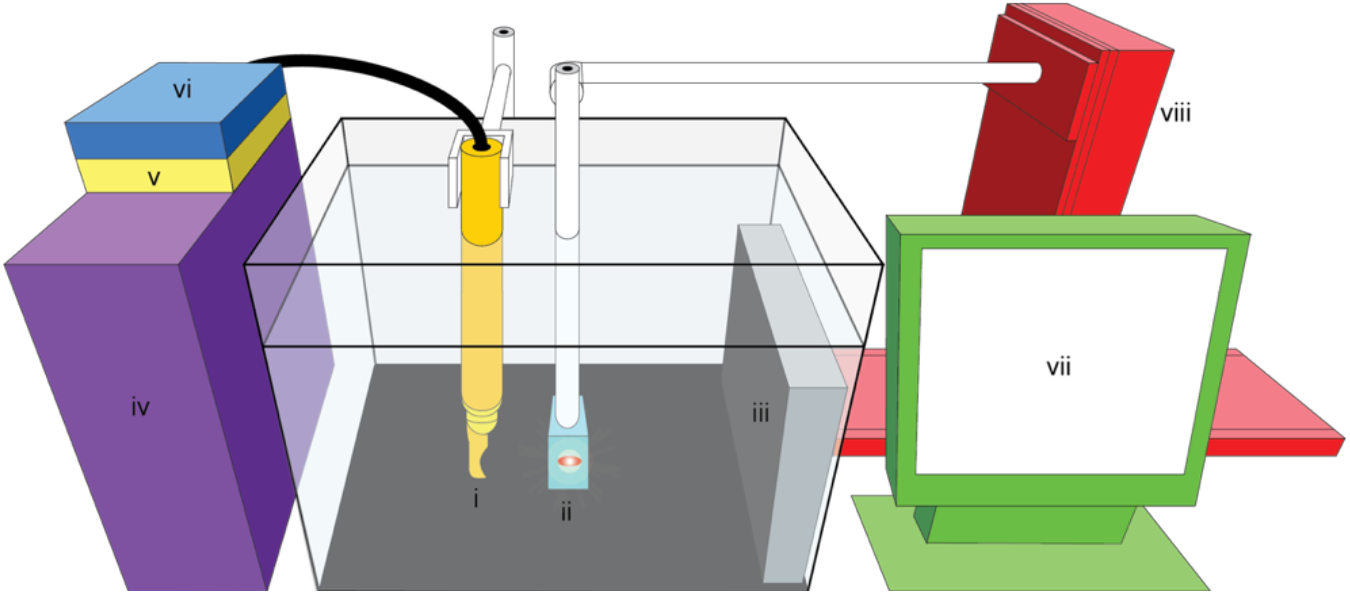
Experimental setup. (i) ultrasound-guided focused ultrasound prototype, (ii) tissue-mimicking phantom, (iii) acoustic absorber, (iv) Vantage 64 LE system, (v) pre-amplifier, (vi) matching network, (vii) screen for real-time monitoring of ablation, and (viii) stepper motor to raise solid water tissue-mimicking phantoms with each sonification.

A 1.620-MHz sonification waveform with a pulse period of 10ms and 50% duty cycle was generated. The solid water sample was sonicated for 10s. To test the built-in imaging probe of the device, brightness-mode (B mode) images were recorded using the imaging transducer described above.

The solid water sample was raised 5mm parallel to the long shaft of the device and, using the aforementioned parameters, an additional lesion was created. This process was repeated to create 5 lesions (**Fig. 4**). During each ablation, ultrasound images were recorded from the built-in imaging probe of the device. A new solid water block was loaded, and the above-described procedure was repeated; however, a 30s sonification was now performed for 3 trials. Axial, lateral, and elevation dimensions of the lesions were measured using a slide micrometer under microscope.

**Fig. 4.**
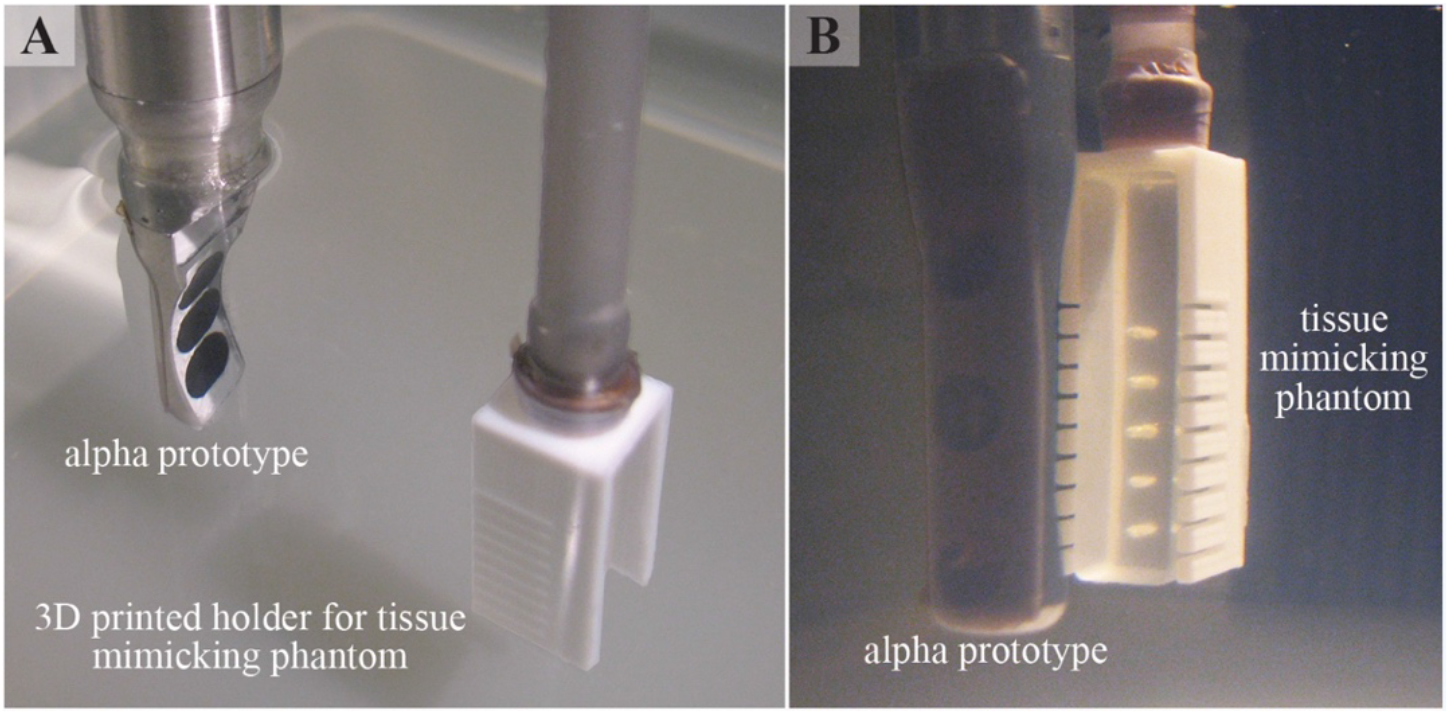
Validation of the prototype. (A) Device aligned with the 3D-printed holder. (B) The prototype generated lesions in tissue-mimicking phantoms secured within the 3D-printed holder.

Subsequently, another block was placed in the water-filled tank at the acoustic maximum, as experimentally detected by a hydrophone (Y-Series High Intensity Hydrophones, Sonic Concepts Inc.). A thermocouple was assembled using a Type T thermocouple connector (SMPW-TM, OMEGA Engineering, Karvina, Czech Republic) and a 40-gauge Type T thermocouple wire with formvar enamel insulation (Pelican Wire, Naples, FL).

The thermocouple was inserted into the block at the acoustic maximum. Temperature measurements were acquired using a microprocessor thermometer (HH23, OMEGA Engineering). Per the aforementioned parameters, a 30s sonification was performed with continuous temperature measurements. As mentioned previously, peak power was applied at 50% duty cycle.

## 3. RESULTS

A prototype device based on Design I (**Fig. 2**) was built, but the design was abandoned due to lateral mode effects that resulted in multiple unwanted focal points and overheating during initial trials. An alpha prototype based on Design II was developed and underwent the above-described testing. At 118W and 50% duty cycle, Design II successfully generated lesions in the tissue-mimicking phantoms (**Fig. 5** & **<u>Supplementary Video 1</u>**). Sonification for a duration of 10s resulted in lesions with average dimensions of 2.5×1.2×6.5mm (lateral, elevation, and axial alignments relative to the transducer). Additionally, lesions created with 30s sonifications resulted in average dimensions of 3.4×3.25×9.36mm (**Table 1**).

**Fig. 5.**
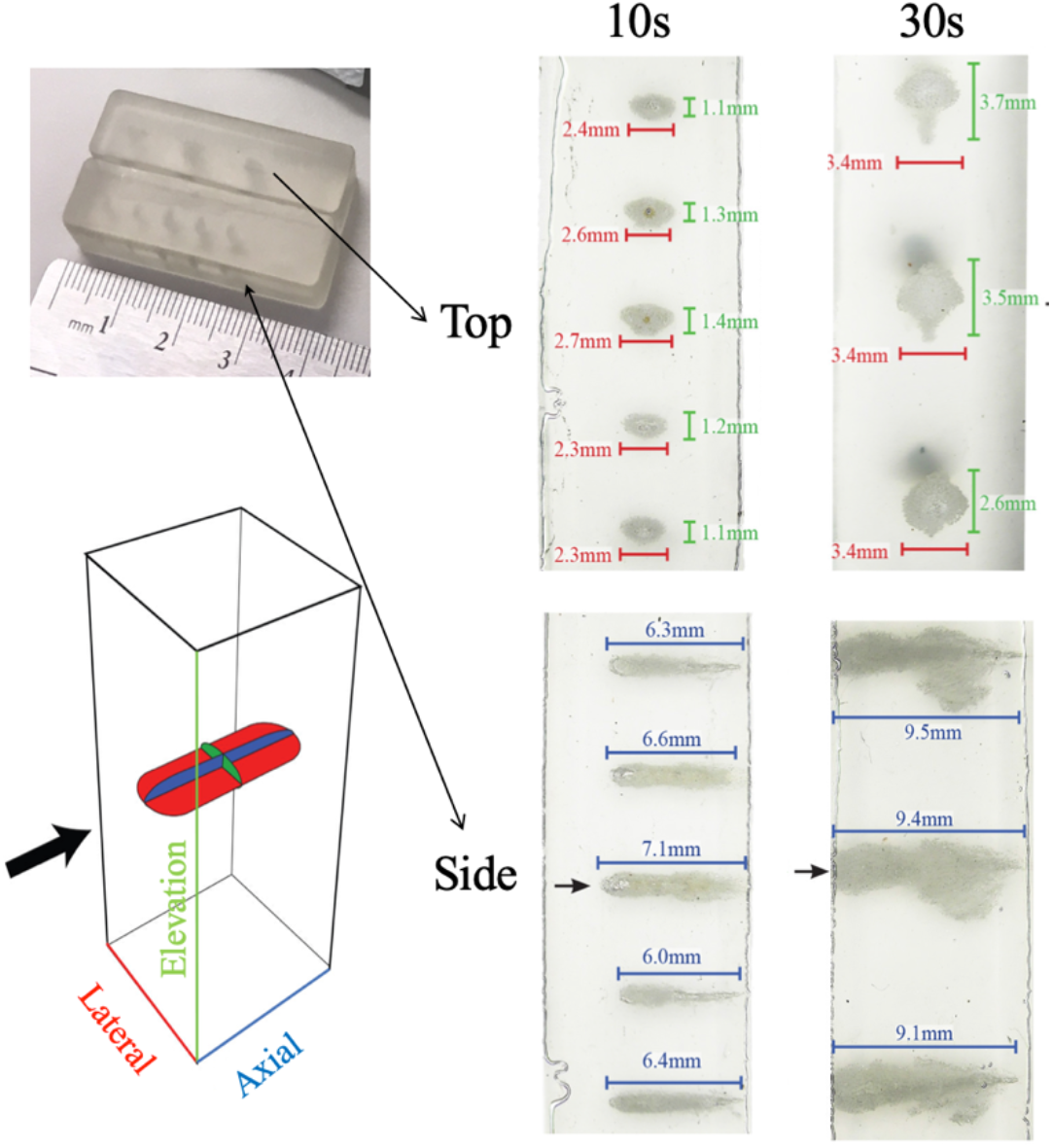
Results of 10- and 30-second ablation of solid water samples. The maximum linear dimension of each lesion created was measured using a slide micrometer under microscope.

**Table 1:**
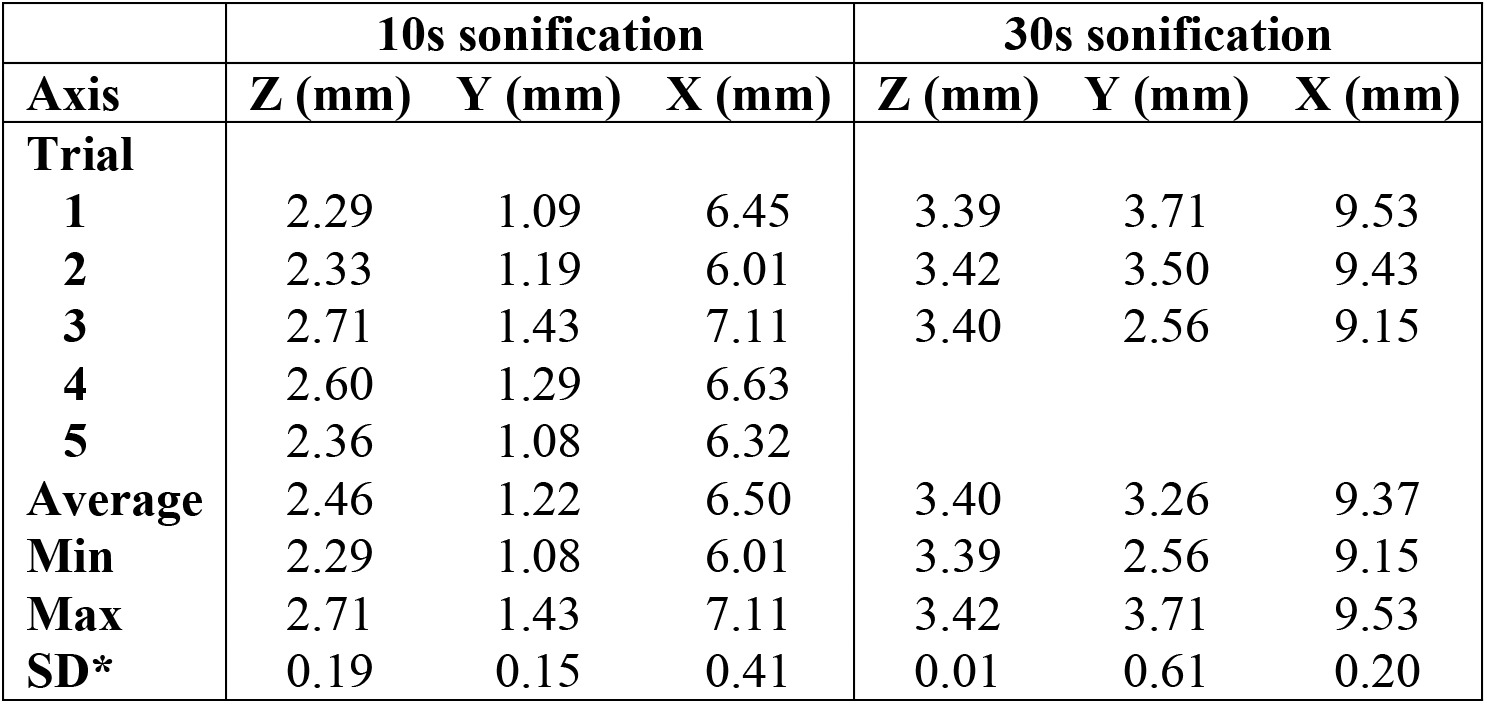
Ablation measurements. *SD, standard deviation.

|  | 10s sonification |  |  | 30s sonification |  |  |
| --- | --- | --- | --- | --- | --- | --- |
| Axis | Z (mm) | Y (mm) | X (mm) | Z (mm) | Y (mm) | X (mm) |
| **Trial** |  |  |  |  |  |  |
| **1** | 2.29 | 1.09 | 6.45 | 3.39 | 3.71 | 9.53 |
| **2** | 2.33 | 1.19 | 6.01 | 3.42 | 3.50 | 9.43 |
| **3** | 2.71 | 1.43 | 7.11 | 3.40 | 2.56 | 9.15 |
| **4** | 2.60 | 1.29 | 6.63 |  |  |  |
| **5** | 2.36 | 1.08 | 6.32 |  |  |  |
| **Average** | 2.46 | 1.22 | 6.50 | 3.40 | 3.26 | 9.37 |
| **Min** | 2.29 | 1.08 | 6.01 | 3.39 | 2.56 | 9.15 |
| **Max** | 2.71 | 1.43 | 7.11 | 3.42 | 3.71 | 9.53 |
| **SD\*** | 0.19 | 0.15 | 0.41 | 0.01 | 0.61 | 0.20 |

Peak temperatures reached 42°C at 10s and 68°C at 30s of sonification. B-mode images acquired before, during, and after ablation resulted in lesion generation (**Fig. 6** & **<u>Supplementary Video 2</u>**). Precise ablation was visualized by the built-in imaging probe during both 10s and 30s sonifications of the phantom blocks.

**Fig. 6.**
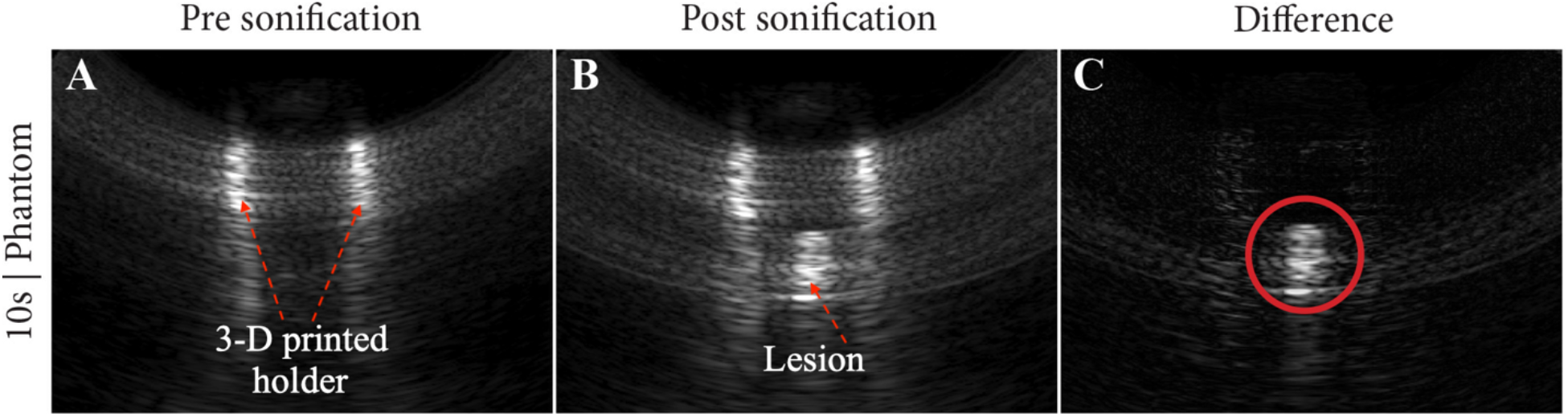
10s sonification of solid water phantom as visualized by the built-in imaging probe: (A) pre ablation, (B) post ablation, and (C) the difference between. The red circle outlines the lesion.

## 4. DISCUSSION

Focused ultrasound devices facilitate targeted tissue ablation from a distance, providing a means to treat pathology previously considered inoperable due to inaccessibility. Noninvasive, transcranial focused ultrasound is limited by the acoustic properties of skull bone.^2,3^ Minimally invasive, intracranial focused ultrasound circumvents these limitations, particularly in patients with excessively dense skulls.^21^ However, developing focused ultrasound devices for use in MIS is challenging. Although acoustic physics favors larger transducer surface areas to achieve ablation, MIS demands tools with increasingly smaller dimensions.^13,14,15^ The optimal geometry for functional, miniature focused ultrasound devices is not well studied.^13^ Therefore, continued size reduction of focused ultrasound transducers requires further investigation of probe acoustic designs (ie, dimensions, curvatures, number of elements, etc.) that can achieve tissue ablation while meeting clinical requirements. This study summarizes a first step in the development of a focused ultrasound device small enough for use in MIS procedures such as neuro-oncological treatment performed with BrainPath™. We have shown that a focused ultrasound device consisting of three 8-mm-diameter circular therapeutic elements arrayed along a 9×32mm cylindrically curved rectangular aperture with 45mm RoC is capable of ablating lesions after 10s or 30s under 1.5MHz (full width half maximum bandwidth: 1.20-1.80MHz)

Previous studies of transcranial focused ultrasound in swine models found histologic evidence of brain tissue coagulation necrosis when focal point temperatures reached 60°C during 12–16s sonification or 55°C during 40s sonification.^22,23^ Calculated thermal dose can be used to predict cell death following focused energy administration.^24,25^ As expressed by the formula for thermal dose 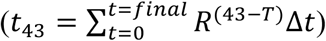, where T is temperature, t is sonification time, and R is related to the temperature dependence of the rate of cell death (R (T < 43°C)=1/4, R (T >43 °C)=1/2), shorter sonification duration with higher thermal rise can achieve comparable thermal dose to longer sonification with lower thermal rise.^24,25^ In the present study, measured focal point temperatures reached 42°C at 10s and 68°C at 30s, suggesting that the presented device can achieve thermal ablation in brain tissue.^23^ Overall, these results indicate that, with further iterations and miniaturizations, the device presented in this study may catalyze the development of new minimally invasive USgFUS devices, particularly those designed for use in neuro-oncological settings.

Finally, the built-in imaging probe of this device offered real-time visualization of the ablation, reducing the need for intraoperative MRI guidance. With a 5MHz center frequency, the image resolution was deemed suitable (based on feedback from our clinical team) for identification of lesions on the tissue-mimicking phantoms; however, for real patients with vasculatures and other complex adjacent anatomies, the resolution of the image guidance will most likely need to be enhanced.

## 5. LIMITATIONS AND FUTURE DIRECTIONS

The device reported in this study was validated using tissue-mimicking phantoms within a controlled test tank, left in 23°C water. However, the thermal dose needed to achieve ablation is known to vary by tissue type, and possessing 37°C.^25^ As a result, additional studies using fresh human brain tissue with histologic evaluation are needed to confirm successful focal point tissue necrosis and the effect on surrounding cells.

Moreover, further *in vivo* testing is required to examine the effects of cerebrospinal fluid and blood flow perfusion on focal point temperatures and surrounding tissue heating. Future studies involving a larger number of array elements can also investigate electronic delays as means to replace mechanical focusing of the probe to target lesions in 3D space. Finally, for the device to be used from within the BrainPath™, additional studies are needed to evaluate ultrasound beam refraction at the interface of the tube and the tissue.

## 6. CONCLUSION

This study reports the development and validation of a USgFUS probe for use in MIS procedures. The dimensional constraints of the prototype were designed to reflect those of BrainPath™ trocars, which are MIS tools used to create atraumatic access to deep-seated brain pathologies. Laboratory testing demonstrated that the MIS USgFUS prototype successfully created lesions in tissue-mimicking phantoms and surpassed threshold temperatures for therapeutic applications. Real-time visualization was also achieved with a built-in imaging probe. Although this study demonstrates successful *in vitro* proof-of-principle, future studies should explore cadaveric validation and additional probe miniaturization for use in MIS procedures.

## Acknowledgments

The authors would like to thank Sonic Concepts, Inc. (Bothell, Washington) for manufacturing the custom probe, as well as Maryland Development Center and Dr. Gil Blankenship for support in software development. NICO Corporation’s Joe Mark and Michele Kennedy are acknowledged for providing their minimally invasive BrainPath™ product for free.

---

**Supplementary Video 1.**
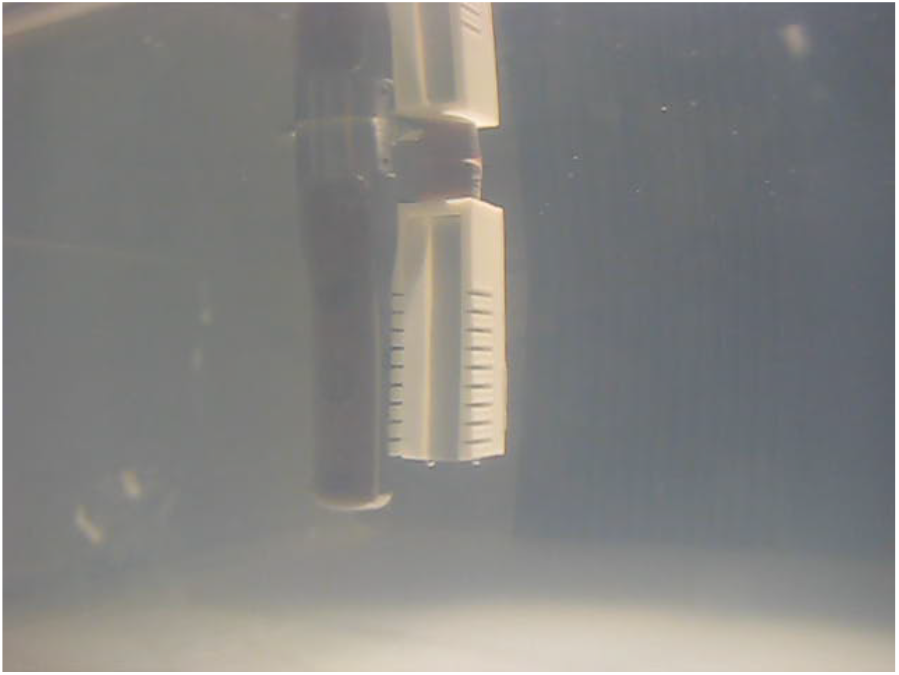
The video demonstrates the reported probe generating lesions in tissue-mimicking phantoms secured within a 3D-printed holder.

**Supplementary Video 2.**
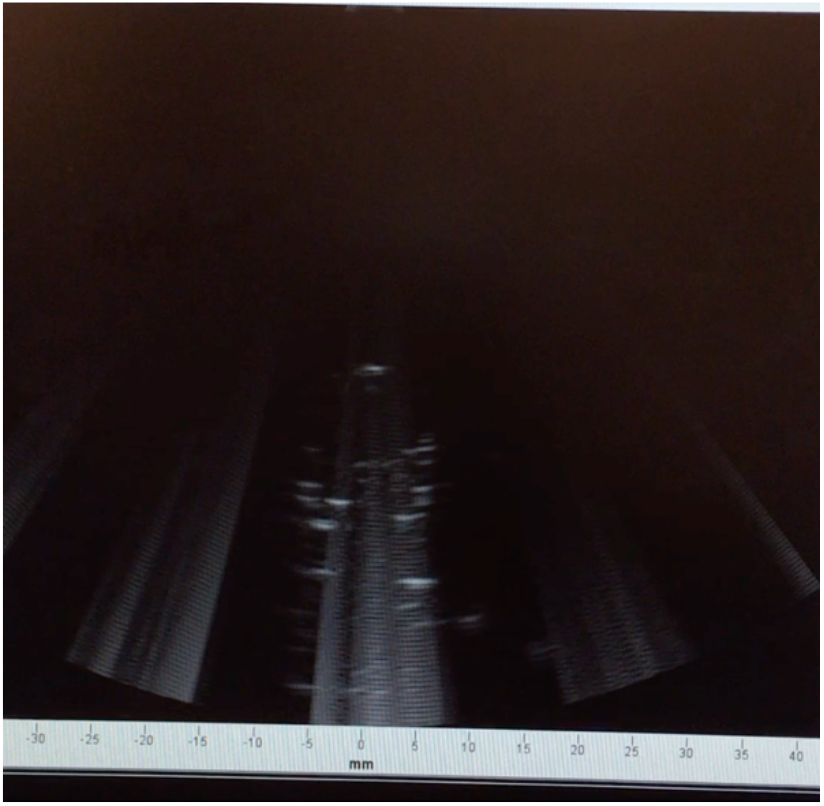
The video demonstrates the real-time ultrasound guidance provided by the built-in imaging probe in the device reported on this study.

